# Deciphering of Somatic Mutational Signatures of Cancer

**DOI:** 10.1101/2022.03.01.482591

**Authors:** Xiangwen Ji, Edwin Wang, Qinghua Cui

## Abstract

Somatic mutational signatures (MSs) identified by genome sequencing play important roles in exploring the cause and development of cancer. Thus far, many such signatures have been identified, and some of them do imply causes of cancer. However, a major bottleneck is that we do not know the potential meanings (i.e., cancer causal or biological functions) and contributing genes for most of them. Here we presented a computational framework, Gene Somatic Genome Pattern (GSGP), which can decipher the molecular mechanisms of the MSs. More importantly, it is the first time, GSGP is able to process MSs from RNA sequencing, which greatly extended the applications of both MS analysis and RNA sequencing. As a result, GSGP analysis matches consistently with previous reports and identify the aetiologies for a number of novel signatures. Notably, we applied GSGP to RNA sequencing data and revealed an RNA-derived MS involved in deficient DNA mismatch repair (dMMR) and microsatellite instability (MSI) in colorectal cancer (CRC).

## Introduction

Understanding of causal and biological processes leading to cancer is important for cancer prevention and drug development. As more and more genome sequencing of tumors have been conducted, cancer genome information allows us to get critical insights from these data, among which the non-negative matrix factorization (NMF)-based somatic mutational signatures (MSs) of cancer derived from these data greatly improved our understanding on the cause and development of cancer^1^.

For example, single base substitutions (SBS) somatic MSs, developed by the Catalogue of Somatic Mutations in Cancer (COSMIC), provided comprehension of various etiological factors of cancer, including genomic deficiencies and environmental exposures. It was reported that SBS2 and SBS13 are attributed to the activity of Apolipoprotein B mRNA editing enzyme catalytic (APOBEC) family^2^, which is the major cause of hypermutation in several cancer types ^3^. SBS3 is proposed to be associated with mutations in BRCA1 and BRCA2 and the homologous recombination DNA repair (HRR) process in which they are involved^4, 5^. SBS4 has been found to be genomic damage caused by smoking and is commonly found in smoking-related cancers such as lung cancer, head and neck squamous carcinoma, and liver cancer^6, 7^. SBS6, together with SBS14, SBS15, SBS20, SBS21, SBS26, SBS44, are known to be associated with defective DNA mismatch repair (dMMR) ^7, 8^. Recognized in cancer patients treated with temozolomide, SBS11 could generate large amounts of mutations^9^. SBS24 is discovered in liver cancer patients with aflatoxin exposure and has been confirmed for causality by experiments using C. elegans^10^ and mice^11^.

Nowadays, MS analysis plays an important role in the field of tumor aetiology, which is used in almost every newly generated tumor genome sequencing study^12, 13, 14^. However, MS analytics has the following bottlenecks: Firstly, although 78 SBS signatures are recorded in COSMIC, the aetiologies of 38 of them are unknown. Moreover, new signatures are still being discovered^15, 16^, posing new challenges to the resolution of signature aetiology. Secondly, the key limitation leading to the previous question is that the signatures revealed by the current studies are complex blends of single base mutation frequency, whose molecular mechanisms remain totally unknown. And third, the current signatures are constructed based on whole-genome sequencing (WGS) or whole-exome sequencing (WES) data, and the use of MS analysis for mutations detected by transcriptome sequencing is an unknown and unexplored field. Moreover, RNA sequencing (RNAseq) derived MSs are being discovered and will confront the first two difficulties as well. Here, we developed an algorithm, Gene Somatic Genome Pattern (GSGP), to decipher the driver genes of MSs. Through case studies, we found that the findings suggested by GSGP demonstrated consistency with previous discoveries. Moreover, by combining the tissue distribution of the signatures, we achieved the positioning of a number of signatures with unknown aetiologies. Strikingly, for the first time, we applied GSGP analysis to RNA sequencing data and predicted and validated an RNA-derived MS related to dMMR and microsatellite instability (MSI) in colorectal cancer (CRC). Finally, an online tool (https://www.cuilab.cn/gsgp) was developed for the personalized GSGP analysis of users. This study provides new insights into deciphering the molecular mechanisms of WES/WGS/RNAseq-based MS analysis, which could yield new understandings of the tumorigenesis and molecular mechanisms of MSs.

## Results

### Assessing genes contributing to the SBS signatures

Briefly, GSGP assigned mutation contexts to genes at or near the mutation sites by weights and solved signatures for each gene using an NMF-based algorithm (Fig. 1a, Methods). Different from the traditional method of calculating signatures for each sample, the GSGP performs signatures for each gene of each sample, which can be used for digging deeper to explore which and how much the signatures/aetiologies affect the genes (or which and how much the genes are affected by the signatures/aetiologies). Specifically, for a sample, GSGP works in 3 steps.

**Fig. 1.**
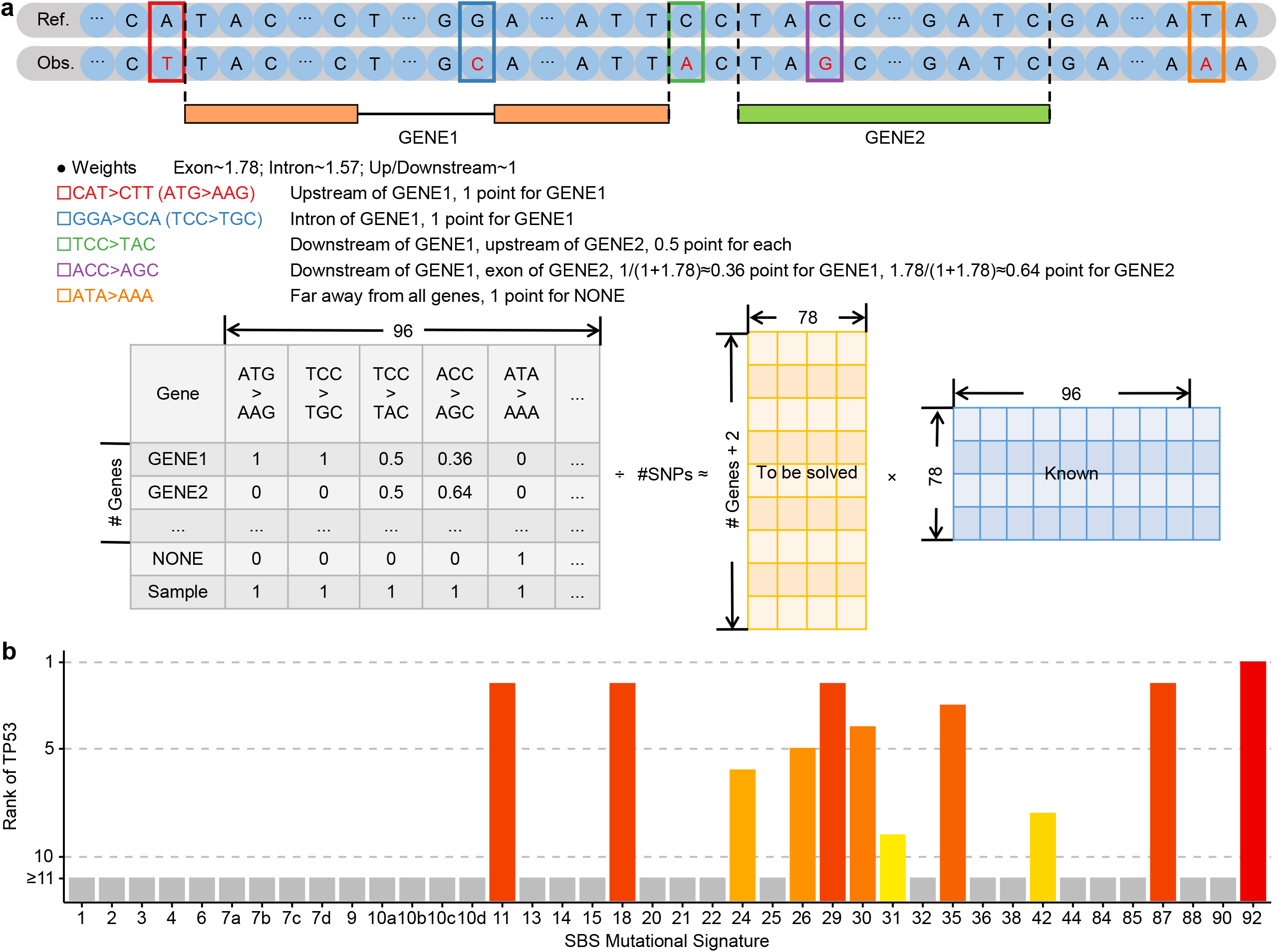
The GSGP workflow. (a) Taking a simulated observation (Obs.), which is aligned to a simulated reference (Ref.) genome and genes (GENE1 and GENE2), as an example, the figure shows how SBS mutation contexts for 5 different conditions are assigned to genes and used to compute MSs, i.e. GSGP. (b) The rank of GSGP of *TP53* among all genes for each signature with known aetiology.

Firstly, we curated the 96 (12 (base replacements) ×4 (5’-type) ×4 (3’-type) =96) different SBS mutation contexts of replacements of pyrimidine (C/T) bases for each mutation site. Same as traditional MS analysis ^1^, the replacements of purines (G/A) can be reverse-complemented to the other strand.

Secondly, the mutation sites are annotated to the exon, intron, upstream, and downstream of genes, so that their corresponding mutation contexts can be assigned to such genes. Particularly, for mutant loci that can be matched to multiple genes, its context is given to each gene with properly-calculated weights according to its position in the mapped genes. A ‘NONE’ gene is introduced when a mutation is far away from all genes.

Finally, given a matrix containing the weighted frequencies calculated previously ((number of genes plus NONE gene) ×96) and a known reference list, we use NMF to decompose and solve the GSGP. Benefiting from the introduction of the ‘NONE’ genes, we can simultaneously solve the sample-level signatures by adding a row vector to the matrix using the sum of each column of it. As a result, we are able to obtain a new dimension of information, genes, for each signature of each sample, compared to the traditional MS analysis.

To test the GSGP algorithm, we applied it to The Cancer Genome Atlas (TCGA) somatic mutation data. As a result, we obtained 78 SBS signatures covering 10,088 samples and 41,924 genes. To confirm the analysis result, we first focused on *TP53*, one of the most critical tumor suppressor genes. The result showed that GSGP ranks *TP53* in the top 10 out of the >40k genes in 11 signatures (Fig. 1b, Supplementary Table 1), which is indeed supported by established knowledge. For instance, SBS24 is the signature associated with aflatoxin exposure and consists mainly of C>A (G>T) mutations. Notably, the G>T mutation in the third position of codon 249 of *TP53* is known to be indicative of aflatoxin exposure^17, 18^. Interestingly, we noticed a couple of treatment-related mutations, including SBS11 (temozolomide), SBS31/35 (platinum), and SBS87 (thiopurine). A *TP53* mutation C380T (S127F), which is the typical base substitution in SBS11, has been reported in the recurrent tumor of a glioma patient treated with temozolomide and was not present in his primary tumor^19^. It is no coincidence that treatments with cisplatin^20^ and thiopurine^21^ have also been shown to induce new mutations in *TP53*. These findings corroborate the possibility of *TP53* mutations caused by these anticancer drugs, which result in the recurrence or progression of cancer.

In addition to focusing on the gene of interest, we can also explore from the signature perspective. SBS4 is widely accepted as a signature caused by smoking. GSGP analysis of this signature indeed revealed a significant contribution of genes related to the sensory perception of chemical stimulus and the behavioral response to nicotine (Supplementary Fig. 1a, 1b). The former mainly includes some olfactory-related receptor genes and the latter is comprised mainly of cholinergic receptor genes, suggesting that smoking could lead to somatic mutations in these genes. Interestingly, a number of genome-wide association studies (GWAS) have revealed that germline polymorphisms in cholinergic receptor genes, including CHRNA3^22, 23, 24, 25^, CHRNA4^26, 27, 28^, CHRNA5^22, 24, 25, 29, 30^, CHRNB1^28, 31^, and CHRNB4^22, 24, 25^, are indeed associated with tobacco addiction and are also associated with the risk of smoking-related diseases including lung cancer. In our previous study, we introduced the cancer germline genome pattern (CGGP) and found a positive correlation between CGGP_E and SBS4^32^. These findings suggest an interaction between genomic variations and environmental exposure, whereby cholinergic receptor germline polymorphisms are associated with tobacco addiction, and tobacco smoking results in somatic mutations in such genes.

### Identification of SBS55 as a new UV related SBS signature

Furthermore, GSGP shows promising potential for exploring the aetiology of signatures. SBS7, which can be separated into 4 subtypes a/b/c/d, has been identified to be involved in DNA damage caused by ultraviolet (UV) exposure^6^. According to the order of wavelength from long to short, UV can be further classified as UVA, UVB, and UVC, among which UVA is the main type that can pass through the atmosphere and cause exposure damage to human^33^. We found that the GSGP profiles of SBS7 subtypes showed significantly different clusters after t-distributed stochastic neighbor embedding (t-SNE)^34^ dimension reduction (Fig. 2a), leading to the different genes and biological functions affected by SBS7 subtypes. Through gene set enrichment analysis^35^ (GSEA), we found that genes contributing to SBS7a were significantly enriched in the ‘response to UVC’ pathway, while genes in SBS7c were identified as ‘response to UVB’ (Fig. 2b), revealing the potential UV wavelength-specificity of UV-related signatures.

**Fig. 2.**
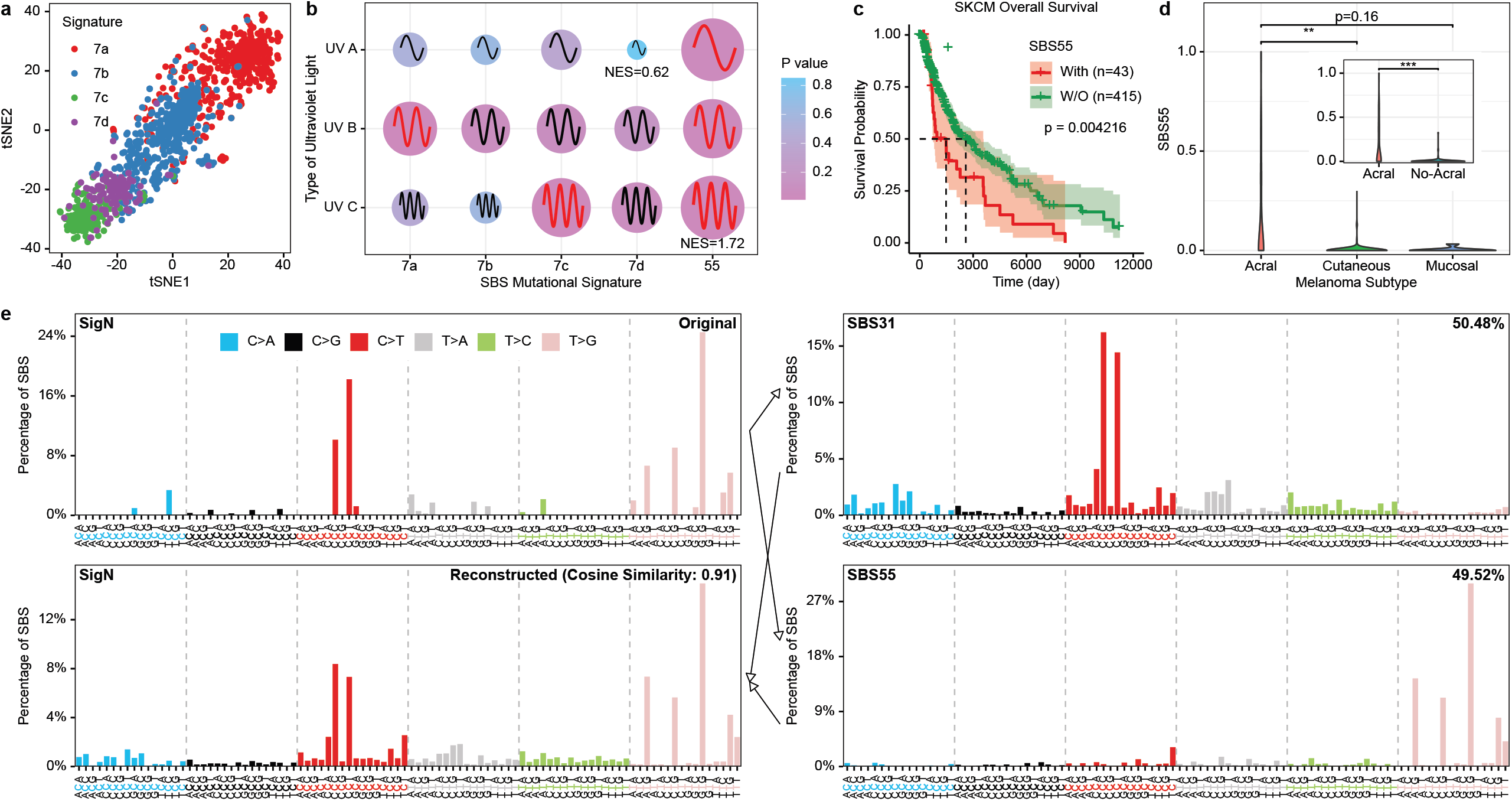
GSGP discovers driver genes and aetiology of UV exposure related signatures. (a) The distinguishability of the GSGP profile of SBS7 subtypes is shown by t-SNE. (b) The ‘response to UV A/B/C’ GSEA of GSGP scores of SBS7a, b, c, d, and SBS55. Point size linearly denotes the normalized enrichment score (NES) and the detailed number of the lowest and highest ones are shown. The waves representing the UV A/B/C (wavelength from long to short) are shown on the points, the waves in red mean significant (p<0.05) results. (c) The Kaplan-Meier curves with 95% confidence intervals show the overall survival of SKCM patients with or without SBS55. (d) The distributions of SBS55 in acral (n=35), cutaneous (n=140), and mucosal (n=8) melanoma. (e) The de novo signature N extracted in TCGA SKCM using SigProfiler can be 49.52% refitted into SBS55.

However, none of the SBS7 subtypes showed significant affection to the genes responding to UVA, the main type causing exposure damage to human^33^. Here, we found that a signature presents only in melanomas, SBS55, was the only signature whose affected genes were enriched in the ‘response to UVA’ process, and the only one that was enriched in all the 3 processes of UVA/B/C simultaneously (Fig. 2b). SBS55 has an unknown aetiology and has been considered a possible signature due to sequencing artefacts. However, by de novo extraction of signatures, we successfully detected the presence of SBS55 in both TCGA and

Australian melanoma biospecimen banks (AMBB) melanoma cohorts^16^, even under a more stringent requirement of read depth (Fig. 2e, Supplementary Table 2), suggesting that it should have some biological significance in melanoma. The results of clustering based on tissue distribution of signatures showed that SBS55 has a closer distance to SBS7 compared to SBS38, which is thought to be non-direct UV damage (Fig. 3a). In conclusion, these analyses revealed that SBS55 represents a deficiency in the defense against UV exposure.

**Fig. 3.**
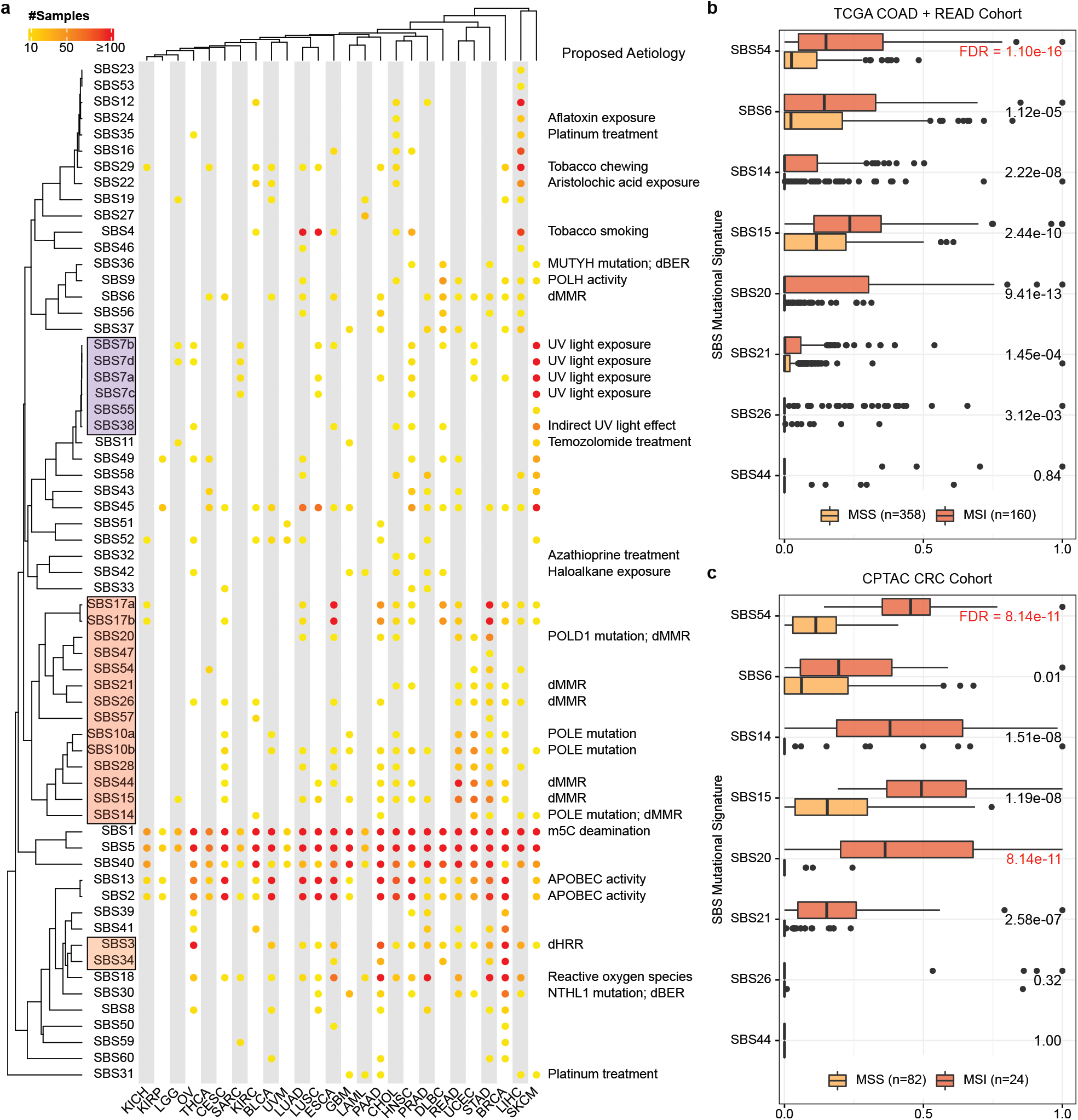
GSGP identifies SBS54 as a new dMMR and MSI related signature. (a) Signatures and cancer types are clustered hierarchically according to the number of samples. The full names of the cancer type abbreviations can be found in https://gdc.cancer.gov/resources-tcga-users/tcga-code-tables/tcga-study-abbreviations. (b, c) The SBS54 shows the strongest ability to distinguish between MSI and MSS CRC samples in (b) TCGA and (c) CPTAC cohorts. Center line, median; box limits, upper and lower quartiles; whiskers, 1.5x interquartile range; point, outliers.

Notably, patients with SBS55 have shorter overall survival days compared to those without (Fig. 2c). Acral melanoma is the melanoma subtype with the most unfavorable prognosis^36^. Because acral melanomas often originate from soles and palms, and their signature profiles are not typically characterized by SBS7, previous studies suggested that acral melanoma is caused by non-UV factors^16, 37^. We found that patients with acral melanoma had significantly higher SBS55 compared with patients with cutaneous or mucosal melanoma (Fig. 2d), hinting that it may be the poorer UV tolerance that leads to the development of melanoma in these areas that are less exposed to UV.

### The positioning of deficient DNA repair associated signatures

In addition to the reasonable result that UV related signatures were clustered into one group by tissue specificity, we observed that a cluster of signatures contains 6 of the 7 known dMMR related signatures (Fig. 3a). There are also 4 DNA polymerase gene mutation-related signatures (SBS10a, SBS10b, SBS14, and SBS20) in this cluster. It is known that DNA polymerases participate in MMR^38^ and base excision repair (BER)^39^. With the study of the genes contributing to the remaining 6 aetiology-unknown signatures (SBS17a, SBS17b, SBS28, SBS47, SBS54, and SBS57) by GSGP, we found that most (5/6) of them are related to MMR and/or BER processes (Supplementary Table 3), suggesting that they could also be involved in dMMR and/or dBER. The dMMR, accompanied with MSI, is frequently observed in CRC^40, 41^, which can be detected by molecular or computational methods^42^. To validate our predictions, we further selected SBS47 and SBS54 which were significantly predicted to be associated with MMR process, while SBS47 was excluded because it was found in only 5 of 518 CRC patients. First, similar to SBS55, we detected the existence of SBS54 in somatic mutation data from 3 independent CRC cohorts using the de novo MS extraction algorithm (Supplementary Table 4). By comparing SBS54 with 7 known dMMR-related signatures, SBS54 had the most remarkable ability to identify patients with microsatellite stability (MSS) and MSI in both TCGA (Fig. 3b) and the Clinical Proteomic Tumor Analysis Consortium (CPTAC) (Fig. 3c) datasets, demonstrating that we successfully discovered a new and potentially more valuable dMMR-related signature using the GSGP algorithm.

SBS3 is the only signature currently proved to be related to dHRR, while SBS34 exhibits a similar tissue distribution (Fig. 3a). The GSGP results show that the genes relevant to SBS34 are enriched in the homologous recombination process (Supplementary Fig. 1c), suggesting that it could be a new dHRR-related signature.

### Towards transcriptomic mutational signature analysis

Transcriptomics is an important branch of tumor omics^43^, and RNAseq can be used for mutation calling as well. However, influenced by complicated transcriptional regulation mechanisms, not all DNA mutations can be transcribed and detected by RNAseq^44^, raising a major challenge for RNAseq-based MS analysis. Besides, the current knowledge of signatures was built based on WGS/WES, which could not be suitable for the signatures decomposed by RNAseq. Here, aiming to investigate dMRR and MSI-related signature in somatic mutations called by RNAseq, we tried to extend the application of GSGP to transcriptome MSs. Firstly, we collected RNAseq data from two CRC cohorts, Chonnam National University and Pusan National University CRC (CP-CRC)^45^ and Zhongnan Hospital CRC (ZN-CRC)^46^, which contain 45 and 72 pairs of tumor-normal paired samples, respectively. When using a priori SBS signatures, we found that neither our identified SBS54 nor the widely established 7 dMMR-related signatures can distinguish between MSI and MSS samples (Fig. 4a, Supplementary Fig. 2), suggesting that brand new de novo extracted signatures need to be characterized in the RNAseq MS analysis. For doing so, we obtained 23 (CP-CRC) and 21 (ZN-CRC) de novo signatures in two datasets and calculated the cosine similarities between the signatures (Fig. 4b), in order to locate the signatures that are stably existing in both datasets. Starting with a classical greedy algorithm, 21 pairs of signatures were matched, of which 7 pairs had a similarity >0.8 (Fig. 4c, Methods). These 7 pairs were then weighted using the sample size and reconstructed into 7 new signatures, named SigA to SigG (Fig. 4d, Supplementary Table 5), in descending order of similarity.

**Fig. 4.**
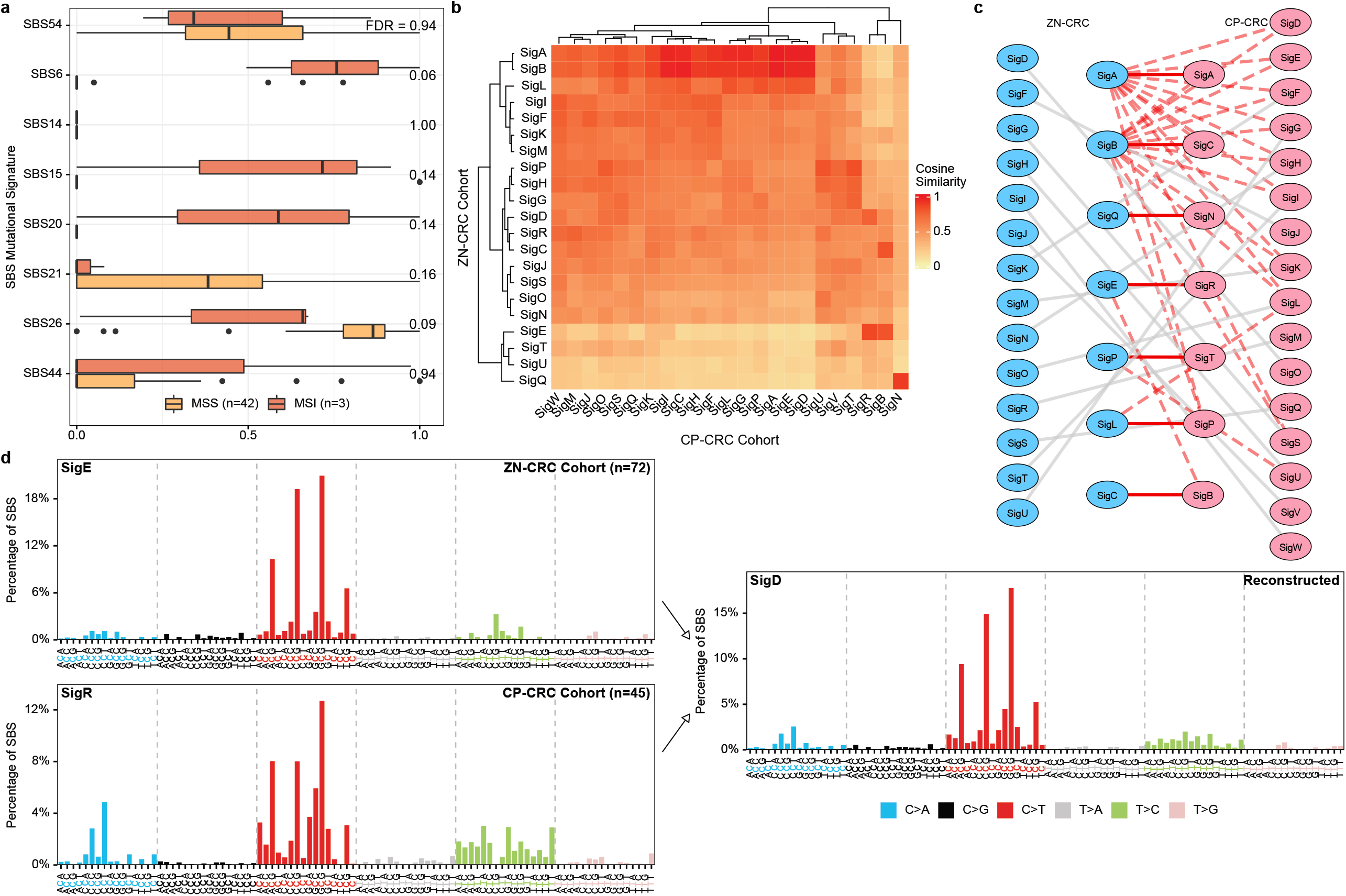
The construction of RNA MSs in CRC. (a) None of the known MSI related COSMIC signatures work in the CP-CRC RNAseq cohort. Center line, median; box limits, upper and lower quartiles; whiskers, 1.5x interquartile range; point, outliers. (b) The cosine similarities between de novo extracted signatures in CP-CRC and ZN-CRC cohorts. (c) The network shows the pairing of signatures. Solid lines are stable pairs while dashed lines are unstable pairs. The line in red means a similarity over 0.8. unstable pairs with similarities<0.8 (should be grey dashed lines) were hidden. (d) Taking SigD as an example, the plot shows the reconstruction of SigD using SigE in the ZN-CRC cohort and SigR in the CP-CRC cohort, weighted by sample sizes.

We next applied GSGP with 7 novel reconstructed signatures to the two datasets to obtain sample-level signatures (Fig. 5a) and GSGP scores of genes. Among them, the GSGP score of SigD is significantly enriched in the MMR process and has the highest normalized enrichment

**Fig. 5.**
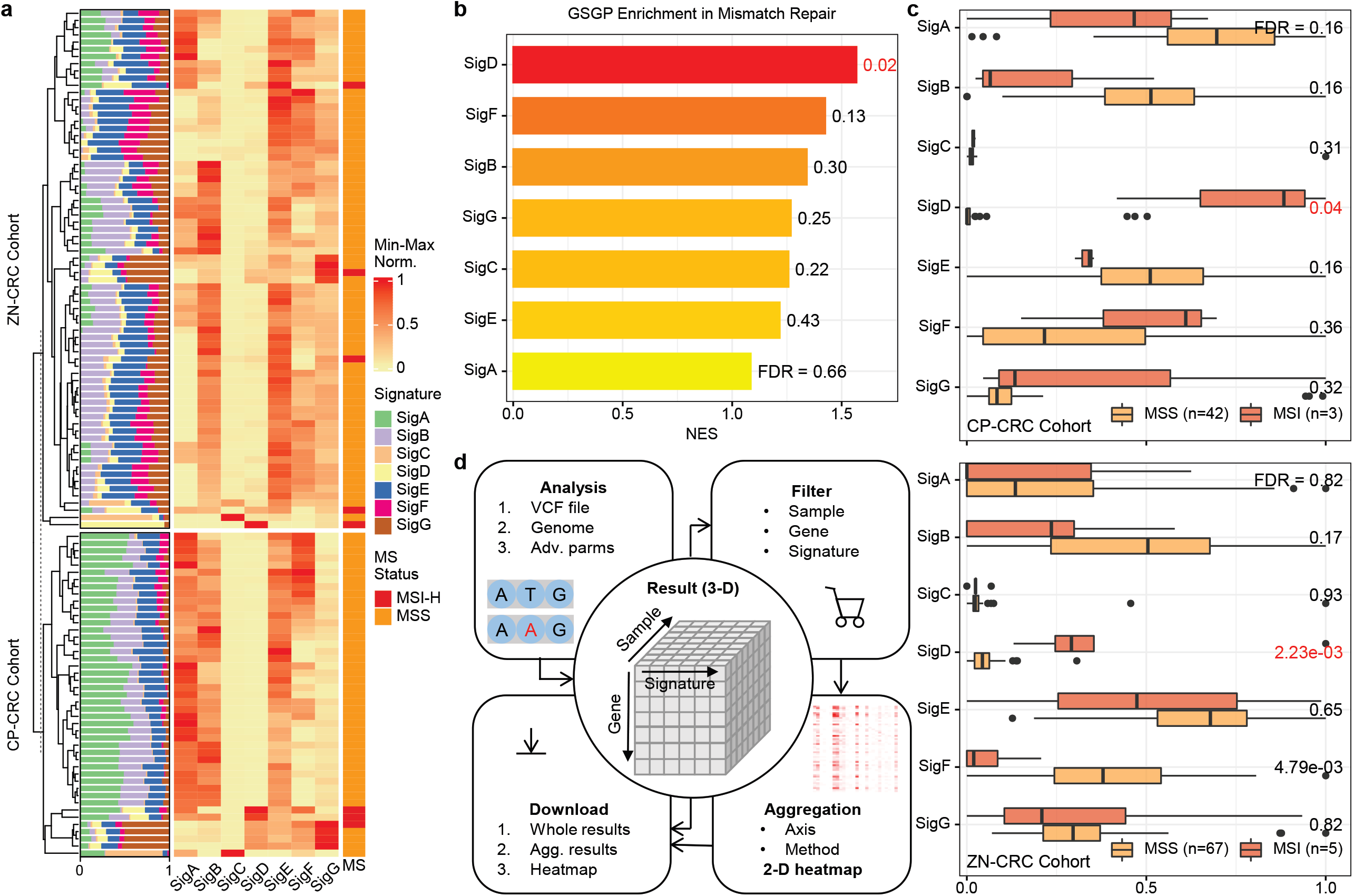
GSGP identifies SigD as a new dMMR and MSI related RNA MS in CRC. (a) The abundance of 7 reconstructed RNA MSs in CP-CRC and ZN-CRC cohorts. The values are min-max normalized by signature and cohort respectively. (b) The mismatch repair gene set enrichment of GSGP score of 7 RNA MSs. (c) The distributions of 7 RNA MSs in MSI and MSS samples in CP-CRC and ZN-CRC cohorts. The most significant results (smallest FDR) are marked in red. Center line, median; box limits, upper and lower quartiles; whiskers, 1.5x interquartile range; point, outliers. (d) The main functions and workflow of the web tool for GSGP analysis. Agg., aggregated.

score (NES) among all signatures (Fig. 5b), implying that SigD is a potential dMMR and MSI-related RNA MS. Furthermore, it is shown that in both cohorts, the patients with MSI have significantly higher SigD compared to those with MSS, while the other signatures do not consistently show such performance (Fig. 5c). These findings indicate that signatures need to be re-identified in RNAseq MS analysis and the GSGP can be used to uncover the signatures with expected aetiology among them.

### The GSGP web tools

Finally, we built a web tool (https://www.cuilab.cn/gsgp) to facilitate the GSGP analysis for researchers (Fig. 5d). The tool accepts the VCF file, one of the most commonly used file formats for storing mutation information, and multiple advanced parameters, and computes 78 SBS GSGPs for each gene of each sample (3-dimensional, 3-D results). For the convenience of visualization, we provide filtering and aggregation functions for the results. Users can specify the axis to be along and method (mean, median, count of values>cutoff) to aggregate the 3-D results into 2-D and display them as a heatmap. The results, as well as a python script that can have more personalized options (customized signatures, species, NMF parameters, multiprocessing, etc.), are available for download.

## Discussion

MS analysis delivers a separation of different cancer somatic mutational processes by an NMF-based method and provides an average reference signature to represent them separately. Current studies have been able to mathematically calculate the mutational burden caused by various mutational processes and experimentally validate the aetiology of some MSs^1^. However, it is still necessary to develop approaches for deciphering mutation patterns to counter the progressively increasing number of signatures. The GSGP algorithm provides a new idea to address this problem. Unlike calculating the frequency of 96 mutation contexts at the sample level, we more precisely assign these mutations to genes and successfully attribute MSs to the gene level. By further joint use of GSEA and tumor distribution similarity, we positioned SBS55 as a new UV exposure-related signature and revealed its deleterious effect on cancer prognosis. Additionally, some signatures involving defective DNA repair including dMMR, dBER, and dHRR were also identified by a similar approach.

As a case study, we investigated the mutational aetiology associated with *TP53*. In addition to common cancer-promoting factors such as smoking^6^ and reactive oxygen species^47^ being associated with *TP53* mutations, which is consistent with existing studies, we found that the use of some cancer therapeutic drugs including platinum, temozolomide, and thiopurine was associated with *TP53* mutations and confirmed the presence of these phenomena in real samples. It is important to note that although each gene showed different levels of contribution to the signatures, these findings do not imply that the mutational process is genetically selective. Another possible explanation is that mutations occurring in other genes caused by the same exposure factors may not play a role in tumorigenesis and therefore would not be observed.

The MS spectrum can be identified by two different methods, the de novo extraction and the reference-based fitting^48^. The former finds a stable decomposition of mutation context patterns by hundreds of runs, which allows both the acquisition of new patterns and the assignment of patterns to existing signatures. The latter uses a reference list of pre-selected known signatures to estimate the mutational burden generated by these mutation processes. In this study, we adopted an approach similar to the latter strategy, that is, we used a reference list consisting of the entire 78 COSMIC SBS signatures for NMF solving. Meanwhile, we used de novo extraction to prove the existence of the signatures we focused on (SBS54 and SBS55). However, the two approaches have their own advantages and disadvantages, respectively^48^. De novo extraction can yield new signatures but may result in excessive merging or splitting of known signatures to the extent that they are undetectable (false negatives). Conversely, reference-based fitting may introduce redundant reference lists and find mutational processes that do not exist (false positives). As a compromise, users can replace their own reference lists to obtain informative results.

Combining germline mutations, somatic mutations, and environmental exposures to construct dynamic models of carcinogenesis is a great challenge^49^. Tobacco smoking is one of the most common environmental carcinogenic factors, which contains a mixture of toxic chemicals that induce DNA damage and are strongly related to the risk of several cancers^50^. Meanwhile, with the accumulation of GWAS studies, there is growing evidence that germline SNPs of some genes show correlations with tobacco addiction in individuals, and some of them are certainly also significantly connected with the risk of multiple diseases caused by smoking, such as lung cancer and COPD. We found that the genes contributing to the smoking-related signature SBS4 include a variety of cholinergic receptors, the main pathway through which nicotine produces pleasure and mediates addiction. However, these mutations are found in tumor tissue and there is no evidence that smoking has mutagenic effects on neurons. A similar idea was proposed by Carmel A. that somatic mutations gained through smoking may be associated with sporadic amyotrophic lateral sclerosis^51^. It has been pointed out that nicotine and cholinergic receptors could participate in the therapeutic resistance of gliomas by activating the stem cell-like phenotype of tumor cells^52^. In addition, nicotine was found to promote brain metastasis from lung cancer by suppressing natural immunity involving microglia^53^. These current studies are focused on smoking-related phenotypes and transcriptomes. More experiments need to be designed to demonstrate the mutagenic effects of smoking on the genome in brain neurons, which could be a novel explanation for the mechanisms of tobacco addiction.

As an important field of omics studies, RNAseq data has been abundantly accumulated^43^. Although RNAseq also allows mutation calling, many mutations detected by WGS/WES cannot be reproduced in RNAseq due to gene expression regulation, such as tissue specificity and alternative splicing^44^. Hence for RNAseq MS analysis, an unknown application field, the knowledge of signatures and their aetiologies that we identified before may no longer be applicable. In this paper, we found that in RNAseq MS analysis, all known dMMR and MSI-related signatures were not effective in differentiating the microsatellite status of patients. Nevertheless, through de novo extraction and reconstruction of signatures, we predicted from 7 reconstructed signatures the potential dMMR and MSI related one, SigD, using GSGP, and proved the accuracy of the prediction in 2 cohorts. Our result is an elegant application of GSGP against these aetiology-unknown signatures. With the wider application in tumor WES/WGS and the expanded application in RNAseq of MS analysis, more and more new signatures will be discovered and urgently needed for aetiology and molecular mechanism explanations, thus the GSGP will be increasingly helpful.

Overall, we presented an efficient algorithm, GSGP, for exploring the driver genes contributing to somatic signatures, a new challenge in cancer genomics. We confirmed the power of GSGP by investigating the *TP53*-contributed signatures and the signature SBS4-driver genes. More importantly, we showed that GSGP is able to explore aetiology for signatures. We revealed that SBS55, SBS54, and SBS34 could be a new UV exposure-related signature, dMMR and MSI related signature, and dHRR related signature, respectively. Even in the novel field of MS analysis for RNAseq, GSGP has demonstrated a surprising prediction performance, i. e., the successful recognition of dMMR and MSI related RNA MS in CRC. In addition, a user-friendly online tool for GSGP analysis was developed. This study provides new insights into deciphering the molecular mechanisms of MSs, which are critical for applying MSs to the development of new therapeutic strategies for cancer.

## Methods

### The gene somatic genome pattern

The workflow of the GSGP algorithm is illustrated in Fig. 1a. Specifically, the calculation of GSGP for a sample can be divided into 3 steps as follows.

Step 1: Like the SBS MS analysis^1^, we obtained the bases of mutation sites and one base at each of their 5’ and 3’ sides, and identified the 96 different mutation contexts of replacements of pyrimidine (C/T) bases. The replacements of purines (G/A) can be reverse-complemented to the other strand. The reference genome FASTA files (hg38, used by TCGA and hg19, used by AMBB) acquired by this step were obtained from UCSC Genome Browser^54^ (http://genome.ucsc.edu).

Step 2: The mutation contexts of each mutation site are assigned to the genes by weights. We used bedtools^55^ (v2.30.0) and its python wrapper, pybedtools^56^ (v0.8.2), to annotate the mutation sites to the exon, intron, upstream, and downstream of genes. For mutations that are at or near only one gene, its mutation context can be assigned to that gene entirely. Yet for mutation locus that can be matched to multiple genes, its context is assigned to each gene by weights according to its position in the gene it is matched to. Based on the research on expression quantitative trait locus (eQTL), we chose appropriate weights, which are exon: intron: up/downstream = 1.78: 1.57: 1, and distance thresholds, which are 121925bp upstream and 121388bp downstream, as described in the section ‘Selection of weights and up/downstream distances’. Particularly, if a mutation locus is far from all genes, it will be assigned to a ‘NONE’ gene. The positions and biotype annotations of genes were collected from Ensembl^57^ (https://www.ensembl.org), and pseudogenes were removed.

Step 3: NMF is used to solve the GSGP. For the weighted counts of mutation contexts of genes counted in the previous step, we first convert them into frequencies by dividing them by the total number of mutations, denoted as matrix X. The rows of X represent genes (including ‘NONE’ gene) and the columns represent the 96 mutation contexts. We solve W, the SBS signature of genes by the following equation:

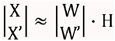

where H (78*96) records the frequencies of 96 mutation contexts occurring in each of the 78 signatures, respectively, which can be downloaded from COSMIC (https://cancer.sanger.ac.uk/signatures/downloads) depending on the different reference genomes. Keeping H unchanged, we solved the GSGP values W using the NMF algorithm from scikit-learn (v1.0.1), whose rows denote genes and columns denote 78 novel SBS signatures. Conveniently, benefiting from the introduction of the ‘NONE’ gene, we can solve for the sample-level signature W’ simultaneously by adding a row vector X’ to X using the sum of each column of X.

### Selection of weights and up/downstream distances

We used a similar strategy to MAGENTA, which is used to perform enrichment analysis of GWAS data^58^, to select thresholds for upstream and downstream distances. Initially, all eQTL data were obtained from the Genotype-Tissue Expression (GTEx) project^59^ v8 (https://gtexportal.org/home). Based on the gene position annotations of GENCODE^60^ v26, which is the version used by GTEx, we annotated eQTL to the exon, intron, upstream, and downstream of the eQTL-correlated gene (eGene). Duplicate eQTL-eGene pairs are kept only once. We used the densities of eQTLs in each case as the weights of mutations. For mutations located on exons, the weights W_exon_ are:

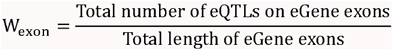

For mutations located on introns, the weights W_intron_ are:

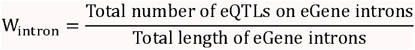

For the case of upstream/downstream, a distance from the gene start/end site was calculated at the same time. We used the median of the distances from all upstream/downstream eQTLs to their eGenes as the threshold for upstream/downstream distances used in GSGP, denoted as D_up_ and D_down_. the weights for mutations within this threshold, W_up/downstream_, are:

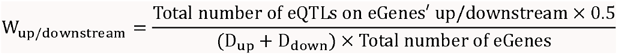

When scaling W_up/downstream_ to 1, the ratio of weights of a mutation located at exon, intron, and upstream/downstream is about 1.78: 1.57: 1 (in detail, 1.78074309122: 1.57446085424: 1).

### Somatic mutation data collection

The somatic mutations of 10,114 samples of 33 different tumor types of TCGA project, as well as their corresponding clinical information, were downloaded from the GDC data portal (https://portal.gdc.cancer.gov). The mutations were called by VarScan2 and stored in Mutation Annotation Format (MAF). We used the ‘maf2vcf’ perl script (https://github.com/mskcc/vcf2maf) to convert the MAF to VCF files. In addition to the filtration implemented by TCGA, the mutations were further filtered using these criteria: (1) only SNVs were used in the SBS signatures we mainly focused on; (2) the read depths covering the mutated sites should be larger than 10; (3) the variant allele frequency (VAF) should be larger than 1%, consistent with our previous research. As a result, 10,088 samples were retained for subsequent analysis.

### RNAseq somatic mutation calling

Two CRC RNAseq datasets were downloaded from the Sequence Read Archive (SRA) (https://ncbi.nlm.nih.gov/sra) with accession number PRJNA748264^45^ (n=45) and PRJNA658001^46^ (n=72). SRA Toolkit (v3.0.0) was used to split the SRA files into pair-end FASTQ files. We used fastp^61^ (v0.23.2) for quality control and adapter trimming for FASTQ. Next, the pair-end FASTQ files were aligned to the human reference genome hg19 in the Spliced Transcripts Alignment to a Reference (STAR)^62^ (v2.7.10a) two-pass mode, and the sorted BAM files were generated. Following the best practice of the Genome Analysis Toolkit (GATK)^63^ (v4.2.5.0), we performed duplicate removing, overhang region clipping, and base quality score recalibration for the BAM files. Mutect2 was used for somatic mutation calling of tumor-normal paired samples and mutations were further filtered by FilterMutectCalls. Also, the same additional 3-step filtering as TCGA described above was applied before performing MS analysis.

### Signature validation using SigProfiler

We used SigProfiler^1^, the de novo MS extraction tool officially provided by COSMIC, to validate the existence of SBS55 in melanoma. Somatic mutation information for 183 melanoma samples was extracted from the AMBB cohort^16^, and the same additional filtration was performed. Then, the python package SigProfilerExtractor (v1.1.4) was applied to the TCGA Skin Cutaneous Melanoma (SKCM) cohort and the AMBB cohort, respectively. Since the low proportion of patients with SBS55, SBS55 may be masked by major signatures such as SBS7a. We disabled the resample to improve the stability of the model to capture more signatures. We selected the optimal number of signatures by 100 replications of NMF solving and assigned the signatures to COSMIC SBS signature v3.2. We also increased the requirement for read depth to >20x and repeated the same MS extraction to exclude the effect of sequencing artefacts.

To validate the existence of SBS54 in CRC, we used 3 independent cohorts from the colon adenocarcinoma (COAD) and rectum adenocarcinoma (READ) projects of TCGA, the CRC project of CPTAC^64^, and the Chinese CRC (CCRC) project^65^. Filtration of depth and VAF was not applied to the latter 2 cohorts because of the unavailability of such information. The same SigProfiler parameters were used for melanoma. Additionally, the same parameters were applied to variants called from 2 CRC RNAseq datasets. However, the purpose was the de novo extraction of signatures, not the validation of the existence of the COSMIC SBS signature.

### Gene set enrichment analysis

For a given signature, we first picked all samples with the signature (sample level SBS signature>0) out. Next, we counted the number of samples with this signature (GSGP value>0) for each gene, which will serve as the scores of the genes (Supplementary Table 1). The genes that did not have the signature in all samples (i.e., those with score=0) were removed because too many equal values would affect the accuracy of gene set enrichment analysis (GSEA). Gene Ontology (GO) gene sets were curated from the Molecular Signatures Database (MSigDB)^66^ v7.1 (http://www.gsea-msigdb.org/gsea/index.jsp). GSEA was implemented in positive score type mode using the R package ClusterProfiler^67^ (v4.0.5).

### Hierarchical clustering

We clustered signatures or tumor types using the number of samples with each signature in each tumor. Numbers of samples were obtained from the detail page of each signature from COSMIC (https://cancer.sanger.ac.uk/signatures/sbs). Min-max normalization was performed on such numbers by each signature. Using the Euclidean distance as the measure of similarity, the complete linkage method as the agglomeration strategy for branches (default parameters), we computed hierarchical clustering using the core function of R (v4.1.0). Leaves were properly ordered by dendsort^68^ (v0.3.4).

### t-SNE analysis of the SBS7 subtype GSGP profiles

For each subtype of SBS7, samples with corresponding signatures present at the sample level were retained, and their GSGP profiles were then extracted and converted into a scanpy^69^ annotated data matrix. To exclude outliers, we removed observations with less than 200 affected genes, and genes with less than 5 observations. A z-score normalization was performed prior to PCA. The first 10 principal components were used for further t-SNE dimension reduction. We used Multicore-tSNE to accelerate the t-SNE operation, which was considered to obtain a better convergence^70^.

### Microsatellite status identification

The microsatellite status of samples from the TCGA COAD and READ projects can be downloaded from the UCSC Xena data hubs^71^ (https://xenabrowser.net/hub/). The microsatellite status of CPTAC^64^ patients is cataloged in LinkedOmics (http://linkedomics.org/cptac-colon/). We used the latest version of a widely accepted tool, MSIsensor-pro^72^ (v1.2.0), to perform the detection of the microsatellite status of 2 CRC RNAseq datasets. The human reference genome hg19 was used to build the reference list and the software was run in tumor-normal paired mode. As recommended, an MSI score >10 was considered as MSI and otherwise as MSS.

### Pairing and reconstruction of RNA mutational signatures

Cosine similarities were used to assess the similarities between two de novo signatures. Subsequently, a classical greedy algorithm, the stable marriage problem, was used to determine a stable pairing scheme for de novo signatures extracted from two RNAseq datasets. Python implementation of this algorithm can be found in https://github.com/Evan210/stable_pairing. Specifically, the signatures of one dataset are called “requesters” and the others are called “acceptors”. Until all “requesters” already have a pair or have asked all “acceptors” for a pair, in each loop: (1) each unpaired “requester” asks for a pair with the “acceptor” who has the highest similarity and is not asked before; (2) each “acceptor” pairs with the highest similar “requester” and rejects the other “requesters”. It is confirmed that whichever side is the “requester” of the pairing will not change the final outcomes of the pairing. As a result, 21 stable pairs were established and 7 of them had cosine similarities >0.8. According to the similarity in descending order, we reconstructed these 7 pairs into 7 new signatures weighted by sample size, named SigA to SigG, respectively.

### Statistical analysis

Survival analysis was performed using the R package survival (v3.2-13) and survminer (v0.4.9), and the log-rank test with one degree of freedom was used to evaluate the significance of the difference in survival days between the two groups. Wilcoxon’s rank-sum tests (python package scipy v1.8.0) were used to estimate the significance of differences between two groups of continuous values. False discovery rates (FDRs) were calculated using the Benjamini-Hochberg method. No same sample was measured repeatedly.

## Data availability

The mutational signature data that support the findings of this study are available from COSMIC, https://cancer.sanger.ac.uk/signatures/downloads. The tumor somatic mutation data that support the findings of this study are available from TCGA, https://portal.gdc.cancer.gov; CPTAC, http://linkedomics.org/cptac-colon/; CCRC, https://doi.org/10.1016/j.ccell.2020.08.002. The RNAseq data used for calling somatic mutations are available from SRA, https://ncbi.nlm.nih.gov/sra, under accession number PRJNA748264 and PRJNA658001. All data generated during this study are included in this article and its supplementary information files.

## Code availability

The GSGP scripts are available at https://www.cuilab.cn/gsgp/download. The stable pairing algorithm is available at https://github.com/Evan210/stable_pairing.

## Acknowledgements

This work was supported by the Natural Science Foundation of China (62025102).

## Author contributions

Q.C. and E. W. conceived of the study. X.J. presented the GSGP algorithm and performed the analysis. X.J., Q.C., and E.W. wrote the paper.

## Supplementary information

### Supplementary Figures and Tables

PDF file containing Supplementary Fig. 1-2 and Supplementary Table 2-4. Detailed legends are provided below each figure/table.

**Supplementary Table 1**

Excel file containing the scores (number of samples with GSGP>0) of genes in each SBS signatures, used for GSEA and the case study of *TP53*.

**Supplementary Table 5**

Excel file containing the 7 reconstructed RNA SBS mutational signatures.

