## Supplementary Figures and Tables for "Deciphering of Somatic Mutational Signatures of Cancer"

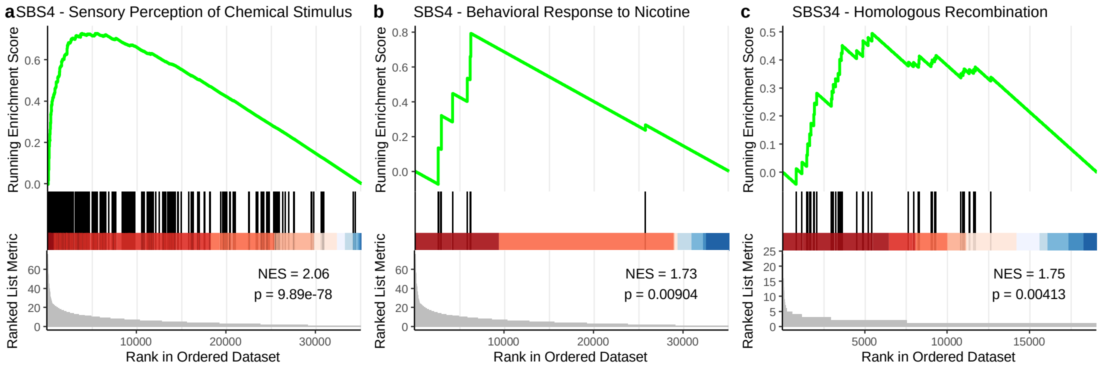


**Supplementary Fig. 1.** The aetiology positioning of signatures using GSEA. The GSEA plot of (a) SBS4 on 'GO: sensory perception of chemical stimulus', (b) SBS4 on 'GO: behavioral response to nicotine', and (c) SBS34 on 'GO: homologous recombination'. NES: normalized enrichment score. The scores used by GSEA are provided in Supplementary Table 1.

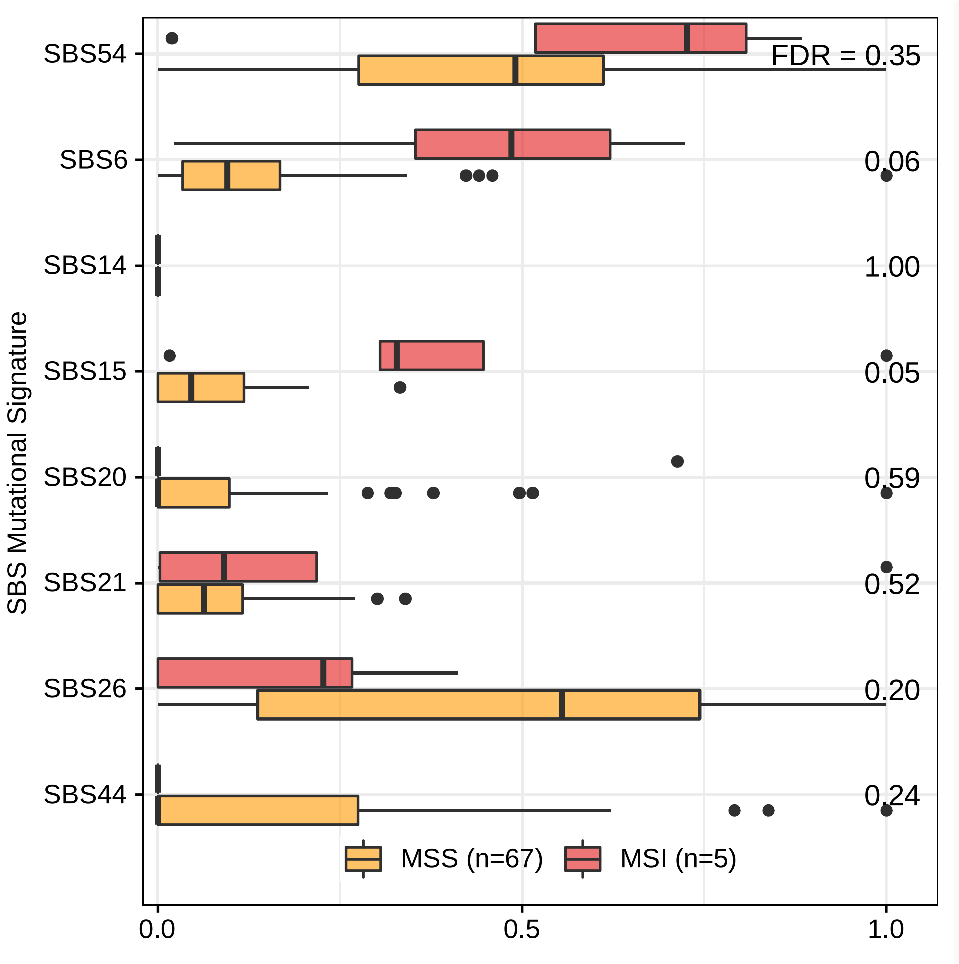


**Supplementary Fig. 2.** None of known MSI related COSMIC signatures work in ZN-CRC RNAseq cohort. Center line, median; box limits, upper and lower quartiles; whiskers, 1.5x interquartile range; point, outliers.

| Cohort | Read depth | Total signatures | Signature containing SBS55 | Assigned signatures | Cosine similarity |
| --- | --- | --- | --- | --- | --- |
| TCGA | >10 | 23 | SigN (14th) | Signature SBS31 (50.48%) & Signature SBS55 (49.52%) | 0.91 |
| AMBB | >10 | 20 | SigM (13th) | Signature SBS31 (38.72%) & Signature SBS55 (61.28%) | 0.87 |
| TCGA | >20 | 22 | SigN (14th) | Signature SBS7a (32.68%) & Signature SBS7b (19.98%) & Signature SBS55 (47.34%) | 0.84 |
| AMBB | >20 | 20 | SigL (12th) | Signature SBS10b (31.26%) & Signature SBS31 (35.56%) & Signature SBS55 (33.18%) | 0.83 |

**Supplementary Table 2.** The de novo signature extractions by SigProfiler prove the presence of SBS55 in melanomas.

| Signature | BER | | MMR | |
|---|---|---|---|---|
| | NES | Pvalue | NES | Pvalue |
| SBS17a | 1.41 | **0.00919** | 1.22 | 0.151 |
| SBS17b | 1.37 | **0.037** | 1.24 | 0.139 |
| SBS47 | 2.28 | **0.0132** | 2.27 | **0.0137** |
| SBS54 | 1.42 | **0.0191** | 1.4 | **0.032** |
| SBS57 | 1.37 | 0.149 | 1.3 | 0.188 |
| SBS28 | 1.45 | **0.0164** | 1.37 | 0.0599 |

**Supplementary Table 3.** The MMR and BER GSEA results of 6 candidate signatures. NES: normalized enrichment score. Significant ($p<0.05$) results were marked in bold.

| Cohort | Read depth | Total signatures | Signature containing SBS55 | Assigned signatures | Cosine similarity |
|---|---|---|---|---|---|
| TCGA | >10 | 23 | SigH (8th) | Signature SBS1 (8.02%) & Signature SBS15 (35.80%) & Signature SBS54 (56.18%) | 0.91 |
| CPTAC | - | 18 | SigF (6th) | Signature SBS1 (5.16%) & Signature SBS20 (53.72%) & Signature SBS54 (41.12%) | 0.88 |
| CCRC | - | 18 | SigH (8th) | Signature SBS15 (71.72%) & Signature SBS54 (28.28%) | 0.81 |

**Supplementary Table 4.** The de novo signature extractions by SigProfiler prove the presence of SBS54 in colorectal cancer.
